# Heterogeneity-driven adaptive scale graph learning for subcellular spatial transcriptomics

**DOI:** 10.64898/2026.05.19.726162

**Authors:** Wanwan Shi, Cong Shen, Ying Liu, Qiu Xiao, Jiawei Luo

**Affiliations:** College of Computer Science and Electronic Engineering, Hunan University, Lushan Southern Road, 410082, Changsha, China; Academy of Mathematics and Systems Science, Chinese Academy of Sciences, 100190, Beijing, China; College of Information Science and Engineering, Hunan Normal University, Lushan Road, 410081, Changsha, China

**Author notes:** Corresponding author: Jiawei Luo:.

**Keywords:** spatial transcriptomics, subcellular resolution, spectral graph filtering

## Abstract

**Motivation:** Spatial transcriptomics enables gene expression profiling within intact tissue sections, providing an important basis for analyzing tissue organization, cellular heterogeneity, and microenvironmental interactions. However, existing spatial structure identification methods often integrate spatial information using fixed neighborhoods or predefined smoothing scales, which limits their ability to adapt to region-specific structural heterogeneity. In homogeneous regions, broader spatial smoothing can help preserve continuous tissue structures, whereas in regions with complex boundaries or mixed cell populations, excessive smoothing may obscure local expression differences and fine-scale structural changes. Therefore, it is necessary to develop an adaptive graph learning framework that can adjust the range of spatial information integration according to tissue structural heterogeneity.

**Results:** In this study, we propose HAST, a heterogeneity-driven adaptive-scale graph learning framework for spatial transcriptomics. HAST adaptively determines graph filtering scales according to spatial structural heterogeneity, enabling flexible information aggregation across different tissue regions. It further decomposes gene expression signals into low-frequency structural components and high-frequency residual components, thereby jointly modeling global spatial continuity and local expression variations. Experiments on high-resolution spatial transcriptomics datasets show that HAST improves spatial structure identification and cross-section generalization. Tumor-enriched cluster identification and neighborhood enrichment analysis further demonstrate its ability to characterize tumor-associated spatial regions and microenvironmental organization.

## Introduction

Single-cell RNA sequencing (scRNA-seq) has revolutionized our understanding of cellular diversity and regulatory dynamics by enabling gene expression profiling at single-cell resolution. However, it inherently loses spatial information during tissue dissociation, hindering the interpretation of cell–cell interactions and tissue organization [1]. Spatial transcriptomics (ST) overcomes this limitation by simultaneously capturing gene expression and spatial context within intact tissue sections. Nevertheless, current ST techniques are limited by their multicellular resolution and cannot resolve molecular distributions within individual cells. Recent advances in spatial molecular imaging technologies have elevated ST to subcellular resolution [2], enabling high-precision mapping of spatial transcriptional profiles and revealing subtle yet critical gene expression differences across subcellular compartments and cell boundaries.

Spatial clustering reveals distinct functional domains and cellular structures, providing valuable insights into cellular functions, heterogeneity, and microenvironmental interactions. Existing spatial clustering methods can be broadly categorized into non-spatial and spatial methods. Non-spatial methods, such as k-means, Scanpy[3], and Seurat[4], rely on gene expression profiles to identify discontinuous domains due to the lack of consideration of spatial information. Researchers have attempted to integrate spatial information to improve spatial domain identification. SpaGCN[5] utilizes graph convolutional networks (GCNs) to identify spatial domains based on the spatial graph. GraphST[6] combines graph neural networks with contrastive learning to enhance the accuracy of spatial domain identification. Spatial-MGCN[7] integrates the spatial and feature graph for comprehensive domain identification. STMIGCL[8] identifies spatial regions by integrating a multi-view graph convolutional network and contrastive learning.

Recently, several studies have applied graph signal processing (GSP) techniques to spatial omics data analysis. SpaGFT[9] introduces a graph Fourier transform framework to decompose spatial expression patterns into different frequency components, providing a new perspective for characterizing spatial structures associated with graph signals at different frequencies. Maher et al.[10] propose a theoretical framework that highlights the biological relevance of high-frequency signals by associating them with intercellular interactions. DeepGFT[11] further integrates graph Fourier transform with deep learning, incorporating spectral information into spatial representation learning. More recently, several studies have explored multi-scale spectral filtering to resolve spatial organization patterns at different hierarchical levels. For example, SpatialZoomer[12] employs a set of low-pass spectral graph filters to extract spatial molecular features across multiple scales, enabling the analysis of spatial organization at the cellular, neighborhood, and tissue-region levels. These studies indicate that spectral filtering and multi-scale information integration provide effective tools for analyzing hierarchical structural patterns in spatial transcriptomics data. However, most existing methods depend on predefined filtering scales or fixed scale combinations, which limits their ability to adaptively adjust the range of information integration according to region-specific structural heterogeneity.

To address these limitations, we propose HAST, a heterogeneity-driven adaptive-scale graph learning framework for spatial transcriptomics. HAST estimates structural heterogeneity from the spatial graph and adaptively adjusts the graph filtering scale, enabling the smoothing range of graph signals to vary according to region-specific structural characteristics. Based on this adaptive scale, HAST performs spectral graph filtering to decompose gene expression signals into low-frequency components that capture global spatial organization and high-frequency residuals that preserve local expression variations. These complementary components are then integrated to learn spatial representations that jointly encode global structural continuity and local heterogeneity. The learned representations are used for downstream analyses, including spatial structure identification and microenvironmental spatial relationship analysis. Extensive experiments based on spatial clustering, cross-section generalization, tumor-enriched cluster identification, and neighborhood enrichment analysis demonstrate the effectiveness of HAST in spatial structure analysis and tumor microenvironment characterization.

## Materials and methods

### Overview of HAST

As shown in Figure 1, HAST consists of three main modules: heterogeneity-driven scale generation, heat-kernel-based frequency decomposition, and graph autoencoder-based representation learning. HAST first constructs a spatial graph based on cell coordinates to model the spatial relationships among cells. To accommodate region-specific structural heterogeneity, HAST estimates graph heterogeneity using edge homophily and adaptively determines the filtering scale for each spatial context. Based on the adaptive scale, heat-kernel spectral filtering is applied to extract low-frequency signals that capture spatially coherent expression patterns, while high-frequency residuals are derived to preserve local expression variations and structural changes. The low-frequency and high-frequency components are then fused to obtain representations that jointly encode global tissue organization and local heterogeneity. Finally, the fused features are fed into a graph autoencoder to learn low-dimensional embeddings through gene expression reconstruction. The resulting embeddings can be used for downstream analyses, including spatial clustering, tumor-enriched cluster identification, and spatial neighborhood enrichment analysis.

**Fig. 1.**
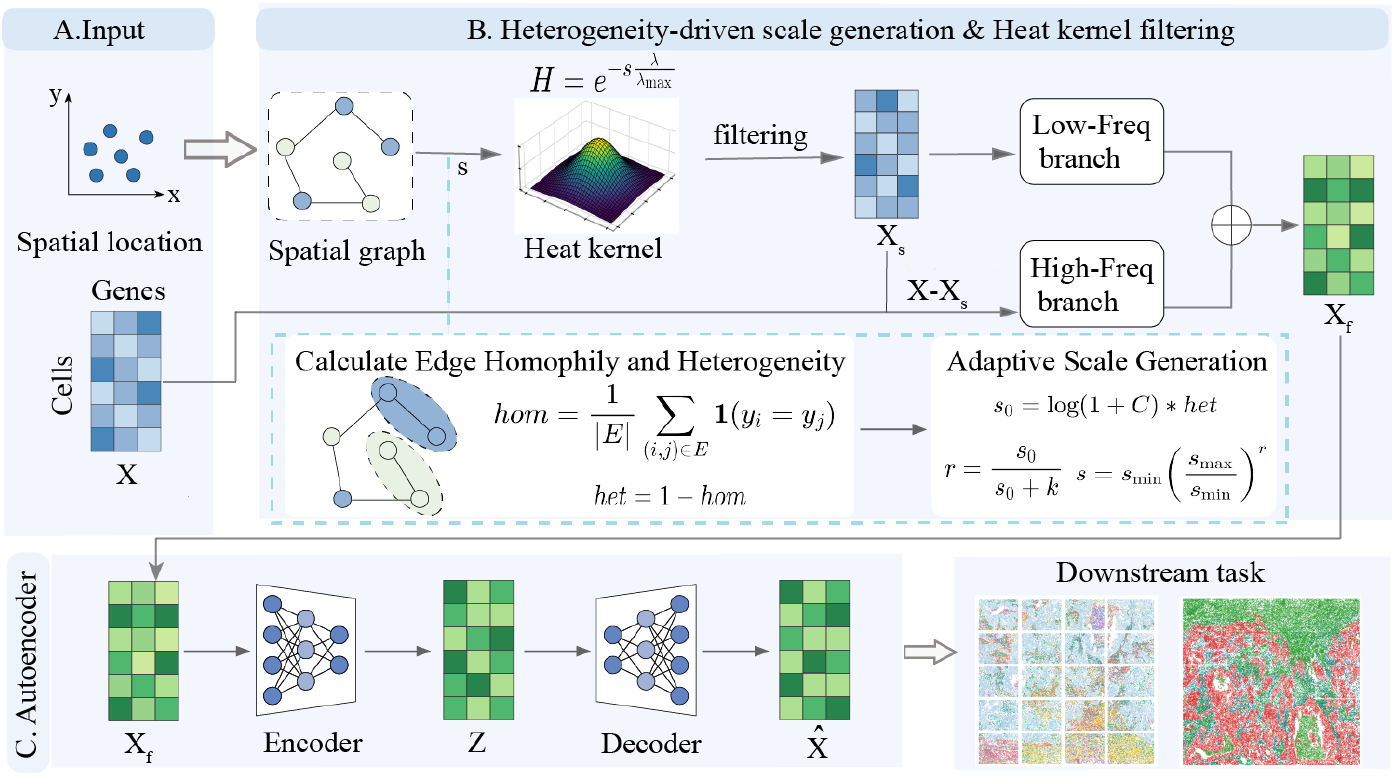
The overall framework of HAST. A HAST takes the gene expression matrix and spatial locations as input. B HAST estimates structural heterogeneity from the spatial graph and adaptively determines the graph filtering scale. Based on this scale, heat-kernel spectral graph filtering extracts low-frequency structural signals, while high-frequency residuals are derived to preserve local expression variations and structural changes. These two components are integrated to generate a fused representation that jointly captures global spatial continuity and local heterogeneity.C The fused representation is encoded into low-dimensional latent embeddings and decoded to reconstruct the gene expression matrix. The learned embeddings are then used for downstream analyses.

### Data preprocessing

HAST takes spatial gene expression data and spatial position information as input. For the spatial gene expression data, HAST employs the SCANPY [3] package to perform log transformation, normalization and standardization by library size on the raw counts. To fully exploit the spatial structure information of spots, we construct an undirected graph *G* = (*V, E*) based on the spatial coordinates of the cell, where nodes represent cells and edges represent connections between cells. Specifically, we compute the Euclidean distance between cells by using their coordinates and build an adjacency matrix *A* ∈ ℝ^*n×n*^, where n denotes the number of cells. If the Euclidean distance between cell i and j is less than a predefined radius r, *A*_*ij*_ = 1; otherwise 0. Considering the differences in sequencing technologies across datasets from different platforms, we set an appropriate radius for each platform. Specifically, the radius *r* is set to 80 for CosMx platform and 500 for osmFISH platform.

### Heterogeneity-driven scale selection

Real tissues often exhibit pronounced spatial structural heterogeneity, where different regions vary in cell composition, structural complexity, and spatial organization [13]. For relatively continuous regions with homogeneous cellular composition, incorporating broader neighborhood information can help preserve spatial continuity. In contrast, for regions with complex boundaries or intermingled cell populations, excessive neighborhood aggregation may lead to over-smoothing and obscure local structural differences [14; 15; 16]. To enable graph signal propagation to adapt to region-specific structural characteristics, we introduce a graph-heterogeneity-driven scale estimation strategy.

Let *y*_*i*_ denote the label of node *i*. The edge homophily of a graph is defined as the proportion of edges connecting nodes with the same label[17]:

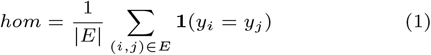

where **1**(·) is the indicator function. A higher edge homophily indicates that adjacent nodes tend to share the same label, suggesting a more homogeneous graph structure. Accordingly, the structural heterogeneity of the graph is defined as:

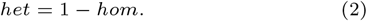

However, graph structural complexity is not determined solely by label differences between neighboring nodes. It is also associated with the diversity of cell types within the tissue [18; 19]. The edge heterogeneity *het* mainly measures whether adjacent cells belong to different categories, and therefore provides a local estimate of structural heterogeneity. When tissues contain different numbers of cell types, graphs with distinct levels of cellular diversity may produce similar edge heterogeneity values. In such cases, edge heterogeneity alone may not fully reflect the increased structural complexity caused by more diverse cellular compositions. To incorporate this effect, we further introduce the number of cell types as a complexity adjustment factor and define the heterogeneity strength as:

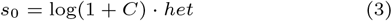

where *C* denotes the number of cell types. The logarithmic transformation log(1 + *C*) is used to retain the influence of cell-type diversity while preventing the scale estimation from being dominated by excessively large values of *C*.

Since the range of *s*_0_ is not fixed, directly using it as the graph filtering scale may introduce unstable scale variations across samples or tissue regions. Therefore, we map the heterogeneity strength into the interval [0, 1) using a nonlinear transformation:

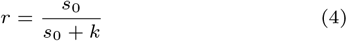

where *k* is a smoothing parameter that controls the sensitivity of the mapping function. This function is monotonically increasing, indicating that a larger heterogeneity strength leads to a larger scale coefficient *r*. Meanwhile, the growth of *r* gradually saturates when *s*_0_ becomes large, which prevents abrupt changes in the estimated scale.

Finally, the scale coefficient *r* is mapped to a predefined scale range [*s*_min_, *s*_max_] to obtain the adaptive graph filtering scale:

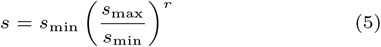

where *s*_min_ and *s*_max_ denote the minimum and maximum allowable scales, respectively. This exponential interpolation provides a bounded and continuous scale estimation strategy. By coupling edge-level heterogeneity with cell-type diversity, the proposed scale selection mechanism enables the graph filtering process to adjust the extent of spatial information integration according to tissue structural heterogeneity, thereby providing an adaptive scale parameter for subsequent spectral graph filtering.

### Spectral graph filtering and feature decomposition

After obtaining the adaptive scale *s*, we further apply spectral graph filtering to the gene expression signal to jointly model global spatial continuity and local structural variations. Let *A* denote the adjacency matrix of the spatial graph, and let *D* be the corresponding degree matrix. The graph Laplacian is defined as *L* = *D − A*, where *D*_*ii*_ = *j A*_*ij*_. A heat kernel filter is adopted to smooth graph signals[20]:

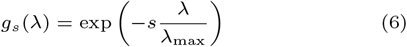

where *λ* denotes an eigenvalue of the graph Laplacian *L*, and reflects the frequency of graph signals in the spectral domain. Smaller eigenvalues are usually associated with low-frequency components that vary smoothly over the graph,whil larger eigenvalues correspond to high-frequency components with rapid local variations. *λ*_max_ denotes the largest eigenvalue of *L*.

The heat kernel filter acts as a low-pass filter on graph signals. It preserves low-frequency components associated with spatially coherent structures while suppressing high-frequency components caused by rapid local changes. By applying the heat kernel filter to the gene expression matrix *X*, the low-frequency smoothed signal can be obtained as:

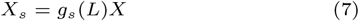

Direct eigendecomposition of the graph Laplacian is computationally expensive, especially for graphs with a large number of nodes. To improve computational efficiency, we approximate the spectral filtering operation using Chebyshev polynomials [21; 22]. Specifically, the filtering operation can be approximated as:

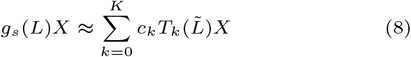

where *c*_*k*_ denotes the filter coefficient, *K* is the polynomial order, 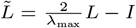 is the rescaled graph Laplacian, and *T*_*k*_(·) denotes the *k*-th order Chebyshev polynomial. The Chebyshev polynomials are recursively defined as:

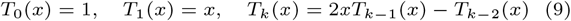

This approximation enables efficient heat kernel filtering without explicitly computing the eigendecomposition of the graph Laplacian.

After deriving the low-frequency structural signal *X*_*s*_, we further extract local variation information from the perspective of frequency decomposition. Since heat kernel filtering mainly retains the low-frequency component, the difference between the original expression signal and the smoothed signal can be regarded as the high-frequency residual. This residual captures local expression differences that are not explained by the low-frequency smoothed signal, including expression changes near structural boundaries and fine-scale spatial variations. The high-frequency residual is defined as:

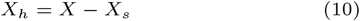

Finally, the low-frequency structural signal and the high-frequency residual signal are integrated to construct the fused feature representation:

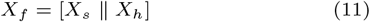

where ∥ denotes feature concatenation along the feature dimension. By combining *X*_*s*_ and *X*_*h*_, the resulting representation *X*_*f*_ retains spatially coherent structural information while preserving local expression variations, thereby providing a more comprehensive representation for subsequent embedding learning.

### Graph autoencoder-based representation learning

Although the fused cell feature matrix *X*_*f*_ integrates both low-frequency structural information and high-frequency residual features, it still lies in a high-dimensional feature space. To learn compact and spatially coherent cell representations, we construct a graph autoencoder framework. Specifically, a graph-attention-based Transformer encoder is employed to model the fused features, thereby learning low-dimensional embeddings that incorporate both spatial neighborhood relationships and gene expression patterns.

For cell node *i*, its representation at the (*l* + 1)-th layer is updated as:

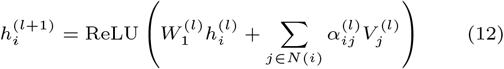

where *𝒩* (*i*) denotes the neighborhood set of node *i*, and 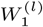 is a learnable weight matrix. The attention coefficient 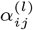 measures the importance of neighboring node *j* to node *i*, and is computed using scaled dot-product attention:

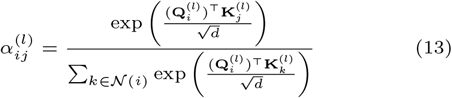

Where, 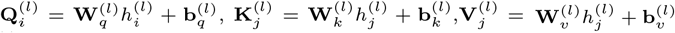, *d* denotes the feature dimension. **Q, K**, and **V** represent the query, key, and value vectors, respectively. 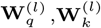, and 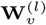 are learnable linear transformation matrices, and 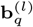, 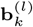, and 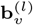 are the corresponding bias terms.

Based on the above graph-attention aggregation scheme, the encoder progressively updates node representations from the fused features *X*_*f*_ and outputs the latent cell embeddings *Z*. Here, *Z* denotes the node embeddings obtained from the last layer of the Transformer encoder. A linear decoder is then applied to map *Z* back to the gene expression space:

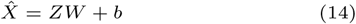

where 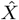 is the reconstructed gene expression matrix, and *W* and *b* are learnable parameters. To ensure that the learned embeddings preserve informative expression patterns, the model is trained by minimizing the reconstruction loss:

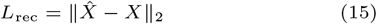

After training, the latent representation *Z* produced by the encoder is used as the low-dimensional cell embedding for downstream analyses.

## Results

### Experimental settings

To assess the performance of our proposed model, we benchmarked HAST against six competing methods on CosMx Lung6, CosMx Lung9 rep1,CosMx Lung9 rep2, CosMx Lung13 and osmFISH mouses omatosensory cortex (osmFISH) dataset[23]. The details of the experimental settings and evaluation metrics can be found in the Supplementary Materials.

### HAST improves the identification of spatial regions

To evaluate the ability of HAST to identify spatial structures, we systematically compared it with six representative spatial clustering methods on the CosMx lung cancer dataset (CosMx Lung9 1). As shown in Figure 2A, HAST effectively recovered the spatial organization patterns of different cell populations, and its clustering results showed a high degree of consistency with the annotated cell types at the overall structural level. Compared with GIST and SIMMT, HAST produced more stable spatial partitions across multiple fields of view (FOVs). Different cell populations formed more compact spatial regions, and the transitions between cell types were more clearly delineated. In particular, in regions where tumor cells were adjacent to fibroblasts and other stromal-associated cells, HAST showed clearer spatial boundaries and better preserved the local distribution patterns of neighboring epithelial cells. This indicates that HAST is able to identify spatial transitions between tumor-associated regions and stromal-associated regions in lung cancer tissues. These results suggest that heterogeneity-driven adaptive scale learning can more effectively capture local structural differences in tissue, thereby improving the recognition of fine-grained spatial distribution patterns.

**Fig. 2.**
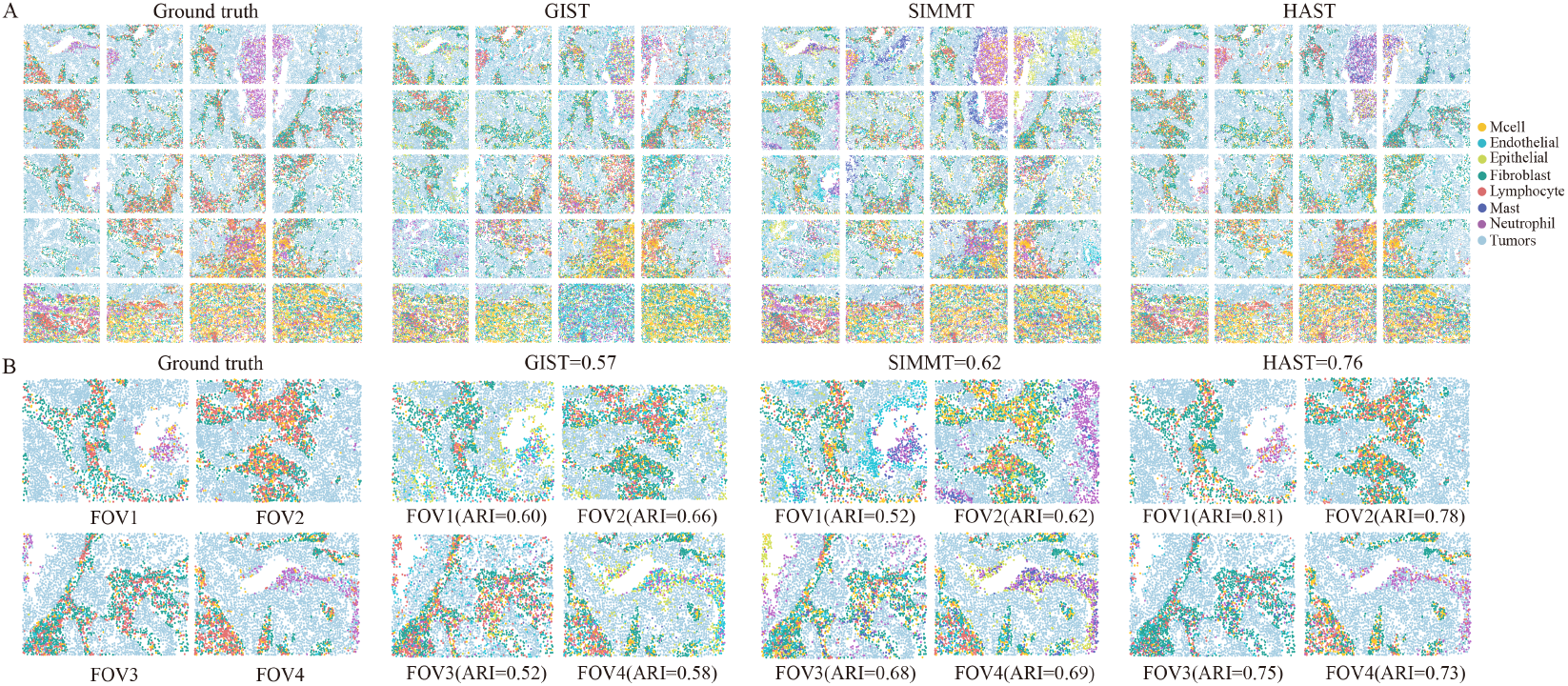
HAST enables accurate identification of spatial regions in the CosMx Lung dataset. (A) Spatial regions from the ground truth and those identified by three different methods across all 20 FOVs of lung cancer tissue. (B) Visualization comparison of representative tumor-dominated FOVs (FOV1: row 3, column 1; FOV3: row 2, column 4; FOV4: row 1, column 1) and a multicellular FOV (FOV2: row 2, column 1).

The overall clustering performance is shown in Figure 2B. HAST achieved the best performance among all compared methods, with an ARI of 0.76. To further evaluate its performance in fine-grained spatial structure analysis, we selected two representative spatial scenarios for visualization, including tumor-dominated FOVs and multicellular FOVs. FOV1, FOV3, and FOV4 represent tumor-dominated regions, where tumor cells occupy a dominant proportion in local areas. In these FOVs, HAST accurately recovered the spatial organization of tumor core regions and clearly distinguished adjacent fibroblasts and immune cells at tumor boundaries. As a result, different cell types formed continuous and stable spatial regions. In contrast, GIST and SIMMT showed cell-type mixing or blurred boundaries in some areas, resulting in less complete spatial structures of tumor regions. FOV2 represents a typical multicellular FOV, which contains tumor cells, fibroblasts, endothelial cells, epithelial cells, and multiple immune cell populations. This region has complex cellular composition and frequent local transitions, making it suitable for assessing the ability of different methods to resolve mixed microenvironments. Compared with GIST and SIMMT, the clustering results produced by HAST were more consistent with the annotated cell types, and different cell populations displayed more natural and continuous spatial distribution patterns. In addition, HAST did not show obvious scattered misclassification or structural fragmentation in the right region of the FOV, suggesting that the model can reduce the influence of noise on spatial structure identification. These results further demonstrate that HAST can maintain global structural stability while preserving local boundary information in complex multicellular microenvironments, thereby more accurately recovering the spatial organization of tissue.

To comprehensively evaluate the generalization ability of HAST across different datasets, we further compared the ARI values of all methods on multiple datasets. As shown in Figure 3, HAST achieved the highest median ARI across all datasets, outperforming the competing methods overall. It should be noted that these datasets contain multiple FOVs, and different FOVs exhibit substantial differences in cellular composition and spatial organization. Despite such structural complexity and heterogeneity, HAST maintained stable clustering performance across datasets. This further indicates that HAST can adapt to diverse spatial structural characteristics and exhibits favorable generalization ability and robustness.

**Fig. 3.**
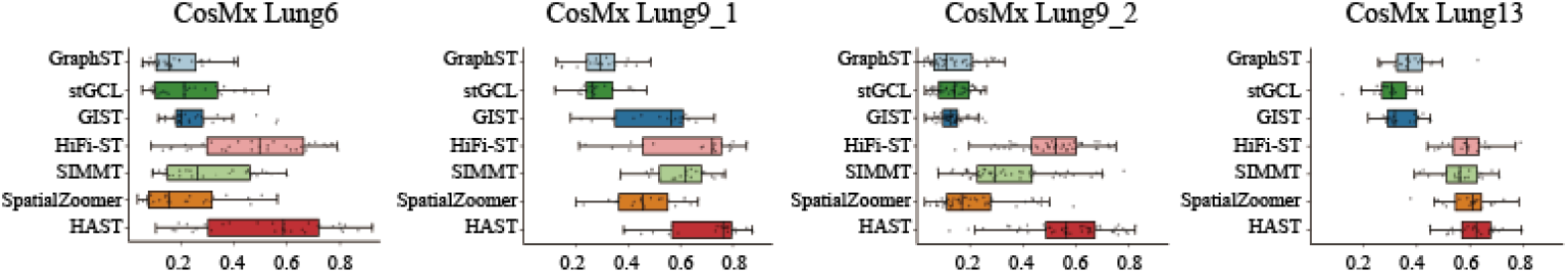
Different ARI of seven methods on the CosMx Lung dataset.

**Fig. 4.**
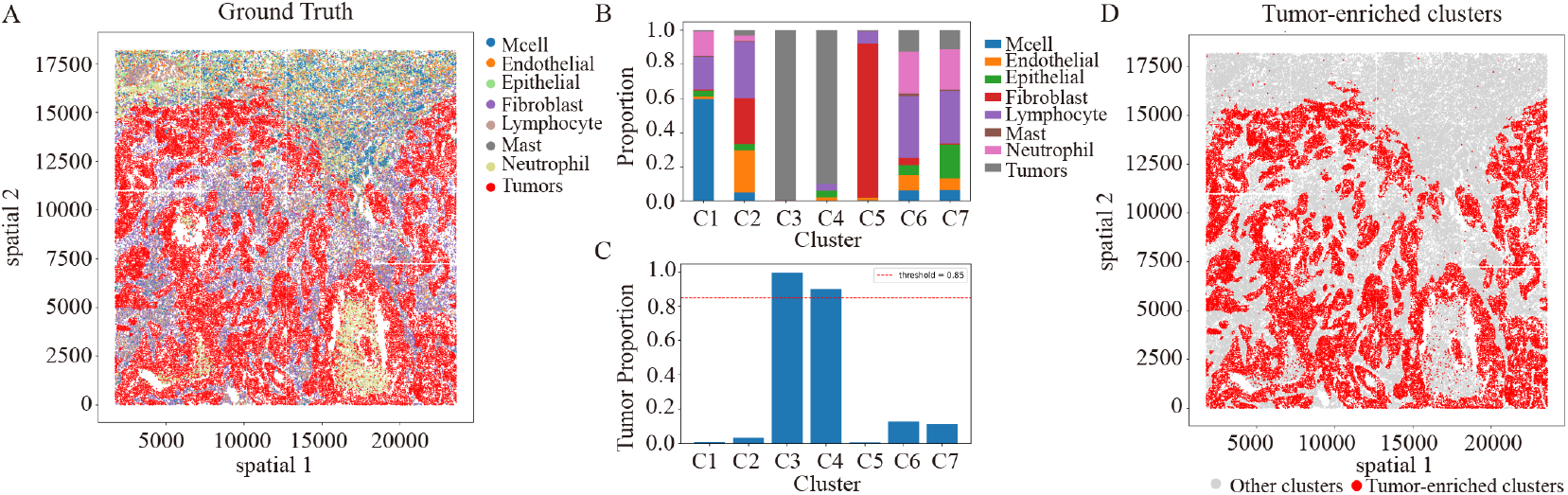
Tumor-enriched cluster identification based on HAST clustering results. (A)Spatial distribution of annotated cell types.(B) Cluster composition analysis showing the cell-type composition of each cluster. (C) Tumor cell proportions across clusters. (D) Spatial distribution of the identified tumor-enriched clusters.

### Identification of tumor-enriched clusters

To further evaluate the ability of HAST to resolve spatial structures in the tumor microenvironment, we performed tumor-enriched cluster analysis on the CosMx Lung9 1 dataset. The dataset contains 20 fields of view (FOVs). To analyze tissue organization at the global spatial scale, all FOVs were first integrated according to their global spatial coordinates. The clustering results were then analyzed within the integrated tissue section. This strategy avoids potential local bias introduced by single-FOV analysis and enables a more comprehensive assessment of the spatial organization patterns of the identified clusters. We first performed cluster composition analysis by calculating the proportions of annotated cell types within each cluster. This step was used to characterize the cellular composition of individual clusters. Based on the proportion of tumor cells in each cluster, tumor-enriched clusters were then identified. Following the definition used in previous studies [24], clusters with a tumor cell proportion greater than 0.85 were defined as tumor-enriched clusters. We further calculated tumor coverage, defined as the proportion of annotated tumor cells contained in the tumor-enriched clusters among all tumor cells. This metric was used to evaluate the extent to which the tumor-enriched clusters covered the spatial distribution of true tumor cells. Spatial visualization was further performed to examine the spatial distribution patterns of the tumor-enriched clusters across the tissue section.

The results are shown in Figure4. The spatial distribution of annotated cell types is shown in Figure4A, where tumor cells formed multiple relatively continuous spatial regions within the tissue. The cluster composition analysis in Figure4B shows that different clusters exhibited distinct cellular compositions, and some clusters were mainly composed of tumor cells. Further quantification of the tumor cell proportion within each cluster showed that clusters C3 and C4 had tumor cell proportions greater than 0.85, and were therefore identified as tumor-enriched clusters Figure4C. As shown in Figure4D, these tumor-enriched clusters formed several continuous spatial regions in the tissue section and were broadly consistent with the spatial distribution of annotated tumor cells, rather than appearing as scattered or random regions. These results indicate that the clustering results produced by HAST can identify tumor-associated spatial regions and effectively reflect the spatial distribution characteristics of tumor cells within the tissue.

## Discussion and Conclusion

Existing spatial structure identification methods usually integrate information from neighboring cells within a fixed spatial range, which limits their ability to adapt to structural differences across tissue regions. To address this limitation, we proposed HAST, a heterogeneity-driven adaptive-scale graph learning framework for spatial transcriptomics. HAST adaptively determines the heat kernel scale according to spatial structural heterogeneity, allowing graph signal smoothing to be adjusted based on tissue organization. Through spectral graph filtering, HAST extracts low-frequency structural signals that reflect global spatial continuity. Meanwhile, high-frequency residual signals are derived from the difference between the original expression signal and the low-frequency smoothed signal to preserve local expression variations. The low-frequency structural features and high-frequency residual features are then integrated and fed into a Transformer encoder to learn cell embeddings that jointly capture global spatial continuity and local heterogeneity.

Comprehensive experiments demonstrated that HAST consistently outperformed six representative baseline methods in spatial clustering across multiple spatial transcriptomics datasets, producing spatial partitions with improved continuity and biological consistency. The tumor-enriched cluster analysis and spatial neighborhood enrichment analysis further showed that HAST could identify tumor-associated spatial regions and characterize the spatial adjacency relationships between tumor regions and surrounding cell populations. These results support the utility of HAST for spatial structure analysis and tumor microenvironment organization. In cross-section generalization experiments, HAST maintained stable predictive performance, indicating that the learned representations preserved structural consistency across tissue sections. Moreover, experiments on the osmFISH mouse somatosensory cortex dataset showed that HAST could recover layer-specific spatial organization under complex anatomical structures and limited gene coverage. Ablation analysis further confirmed that the joint modeling of low-frequency structural information and high-frequency residual information contributed to more informative spatial representation learning.

## Notes

### Competing Interest Statement

The authors have declared no competing interest.

### Funding

- National Natural Science Foundation of China: 62372165

